# A new oomycete metabarcoding method using the *rps10* gene

**DOI:** 10.1101/2021.09.22.460084

**Authors:** Zachary S. L. Foster, Felipe E. Albornoz, Valerie J. Fieland, Meredith M. Larsen, F. Andrew Jones, Brett M. Tyler, Hai D. T. Nguyen, Treena I. Burgess, Carolyn Riddell, Hermann Voglmayr, Frank N. Martin, Niklaus J. Grünwald

**Author notes:** Corresponding authors: Frank Martin, Niklaus Grünwald.

## Abstract

Oomycetes are a group of eukaryotes related to brown algae and diatoms, many of which cause diseases in plants and animals. Improved methods are needed for rapid and accurate characterization of oomycete communities using DNA metabarcoding. We have identified the mitochondrial 40S ribosomal protein S10 gene (*rps10*) as a locus useful for oomycete metabarcoding and provide primers predicted to amplify all oomycetes based on available reference sequences from a wide range of taxa. We evaluated its utility relative to a popular barcode, the internal transcribed spacer 1 (ITS1), by sequencing environmental samples and a mock community using Illumina MiSeq. Amplified sequence variants (ASVs) and operational taxonomic units (OTUs) were identified per community. Both the sequence and predicted taxonomy of ASVs and OTUs were compared to the known composition of the mock community. Both *rps10* and ITS yielded ASVs with sequences matching 21 of the 24 species in the mock community and matching all 24 when allowing for a 1 bp difference. Taxonomic classifications of ASVs included 23 members of the mock community for *rps10* and 17 for ITS1. Sequencing results for the environmental samples suggest the proposed *rps10* locus results in substantially less amplification of non-target organisms than the ITS1 method. The amplified *rps10* region also has higher taxonomic resolution than ITS1, allowing for greater discrimination of closely related species. We present a new website with a searchable *rps10* reference database for species identification and all protocols needed for oomycete metabarcoding. The *rps10* barcode and methods described herein provide an effective tool for metabarcoding oomycetes using short-read sequencing.

**Interpretive summary:** Oomycetes are a group of eukaryotes related to brown algae and diatoms, many of which cause diseases in plants and animals. Improved methods are needed to rapidly characterize the diversity of oomycete species found in environmental samples. We have identified the mitochondrial 40S ribosomal protein S10 gene (*rps10*) as being useful for oomycete community sequencing. We evaluated its utility relative to a popular barcode, the internal transcribed spacer 1 (ITS1), by sequencing environmental samples and a community we synthesized in the laboratory. The amplified *rps10* region is predicted to have a higher taxonomic resolution than ITS1, allowing for greater discrimination of closely related species. We present a new website with a searchable *rps10* reference database for species identification and all protocols needed for oomycete community sequencing. The *rps10* barcode and methods described herein provide an effective tool for characterizing oomycetes using environmental DNA sequencing.

## 1. INTRODUCTION

Oomycetes are microscopic eukaryotes related to brown algae and diatoms that often cause diseases in plants and animals (Baldauf, Roger, Wenk-Siefert, & Doolittle, 2000; Yoon, Hackett, Pinto, & Bhattacharya, 2002). They include highly destructive pathogens with major impacts on agriculture (Fry, 2008), aquaculture (Phillips, Anderson, Robertson, Secombes, & van West, 2008), and natural ecosystems (Cahill, Rookes, Wilson, Gibson, & McDougall, 2008; Grünwald, LeBoldus, & Hamelin, 2019). Oomycetes that cause agricultural diseases include: *Phytophthora infestans*, the cause of potato late blight and the Irish Potato Famine (Fry, 2008); *Aphanomyces euteiches,* the pathogen responsible for damping-off and root rot of legumes (Gaulin, Jacquet, Bottin, & Dumas, 2007); *Pythium* species that cause damping-off and root rot on a large variety of agricultural and horticultural crops (F. N. Martin & Loper, 1999); and host-specific obligate pathogens known as the white rusts and downy mildews (Spring et al., 2018).

In addition to their impacts on agriculture, invasive oomycete pathogens have destructive effects on forests, managed landscapes, and aquatic ecosystems (Hansen, Reeser, & Sutton, 2012). Forests in North America and Europe have suffered significant tree mortality due to sudden oak death and larch death, respectively, caused by *Phytophthora ramorum* (Brasier & Webber, 2010; Grünwald, Goss, & Press, 2008). Eucalypt forests in Australia are experiencing massive dieback caused by invasive *Phytophthora cinnamomi* (Burgess et al., 2017). Tree seedlings in natural ecosystems regularly suffer damping off, which is frequently associated with *Pythium* species (Augspurger & Wilkinson, 2007). Some oomycetes are also significant fish and crustacean pathogens, such as *Saprolegnia parasitica* and *Aphanomyces invadans*, and are of great concern in aquaculture (van West, 2006). Because of the ecological and economic impacts of oomycetes, improving methods to characterize the distribution of these pathogens quickly, accurately, and economically would help control and understand this important, but relatively understudied, group of organisms.

Although many oomycete pathogens are well known for the extensive damage they cause to natural ecosystems and agriculture (Wills, 1993), the global diversity and distribution of oomycetes as a whole is less well characterized compared to other microorganisms, such as fungi and bacteria. This is due, in part, to the challenge of collecting samples during periods when oomycetes are active, isolating them from diverse host and substrate materials (e.g., water, soil, plant, and animal tissues), culturing them on an assortment of specialized media (or *in vivo* for the numerous obligate pathogens), and identifying species using morphological characteristics. Methods relying on DNA sequencing are increasingly used to complement or replace these traditional techniques. Culture-independent DNA-based methods like metabarcoding have the potential to overcome many of the challenges associated with characterizing oomycete communities (Salcedo et al., 2021; Tedersoo, Drenkhan, Anslan, Morales-Rodriguez, & Cleary, 2019), as has been demonstrated with fungi (White et al., 1990; Nilsson et al., 2019; Schoch et al., 2012) and bacteria (Bukin et al., 2019; Tringe & Hugenholtz, 2008). In order for metabarcoding of oomycetes to be effective, a region of DNA ideally should: (1) contain enough variability to differentiate isolates to species level, (2) be small enough for short-read, high-throughput sequencing, and (3) be flanked by regions where primers can be designed that are conserved in oomycetes, but diverged in other organisms (Cristescu, 2014). In addition, a curated database of high-quality reference sequences of the barcode region must be publicly available so environmental sequences produced by metabarcoding can be assigned taxonomic classifications. In practice, it is difficult to find a barcode with all these properties. Many widely used barcodes/primers fail to differentiate closely related species, often amplify non-target organisms, or fail to amplify target organisms.

Reads generated by metabarcoding are generally clustered into operational taxonomic units (OTUs) or amplified sequence variants (ASVs). Both methods serve to mitigate sequencing error by grouping similar sequences. However, OTUs are created by clustering at a specific sequence similarity threshold, usually meant to simulate species-level differences, whereas ASVs are created using a statistical model of mutation and read abundance to correct for sequencing errors (Callahan et al., 2016). OTU clustering is the older and more established of the two methods. The threshold used for OTU clustering is specific to the locus and group of organisms analyzed. This threshold is somewhat arbitrary, considering that different OTU clustering techniques apply this threshold differently (Chen, Zhang, Cheng, Zhang, & Zhao, 2013), but is designed to simulate species-level differences. ASV inference requires no pre-determined clustering threshold, but rather similar sequences are grouped to the extent needed to correct for sequencing errors based on a model of mutation frequencies inferred from the data. Ideally, ASVs represent real biological sequences and can be compared between studies, whereas OTUs are emergent properties of a specific dataset and cannot be compared between studies (Callahan, McMurdie, & Holmes, 2017). ASV inference is a relatively new but currently preferred method due to it robust statistical foundation and ability to produce results that are comparable across studies.

There have been several published DNA barcodes for oomycetes (Choi et al., 2015; Robideau et al., 2011; Yuan, Feng, Zhang, & Zhang, 2017). However, the most popular, the internal transcribed spacer 1 of the ribosomal DNA (ITS1), has insufficient taxonomic resolution to identify many oomycetes to the species level (Redekar, Eberhart, & Parke, 2019), which can lead to ambiguous or incorrect taxonomic classifications (Riddell et al., 2019). Furthermore, currently available ITS1 primers amplify distantly related non-target organisms such as plants and fungi (Coince et al., 2013) or only amplify some oomycete genera (Legeay et al., 2019). A widely used method uses a semi-nested PCR with the primers ITS6 and ITS4 in the first reaction followed by ITS6 and ITS7 in the subsequent reaction (Cooke, Drenth, Duncan, Wagels, & Brasier, 2000). In practice, as little as 5.3% of the OTUs (57% of reads) recovered using this method were assigned to oomycetes (Coince et al., 2013). Sapkota and Nicolaisen (2015) proposed increasing the annealing temperature to increase specificity to oomycetes and reported 60% of OTUs (95% of reads) assigned to oomycetes. Riit et al. (2016) developed primers targeting the ITS1 and ITS2 regions without the need for a semi-nested approach; those primers enabled assignment of 22% and 29% of OTUs to oomycetes (25% and 30% of reads, respectively). Other primers for ITS1 and other loci, including cytochrome c oxidase subunit I (*cox1*) and cytochrome c oxidase subunit II (*cox2*), have been used, but have either not been extensively tested or only target the genus *Phytophthora* (Esmaeili Taheri, Chatterton, Gossen, & McLaren, 2017; Fiore-Donno & Bonkowski, 2021; Landa et al., 2021; Riddell et al., 2019; Sapp et al., 2016). In summary, all the ITS1-based methods targeting oomycetes we are aware of have shortcomings regarding non-target amplification and/or taxonomic resolution.

The mitochondrial *rps10* gene, encoding the 40S ribosomal protein has been helpful in delineating species and estimating phylogenetic relationships in *Phytophthora* (F. N. Martin, Blair, & Coffey, 2014). Preliminary analysis using mitochondrial genomes from a broad range of taxa identified an arrangement of tRNA genes flanking the *rps10* gene unique to oomycetes (*tRNA-Phe, rps10, tRNA-Arg, tRNA-Gln, tRNA-Ile, tRNA-Val*; F. Martin, unpublished). Amplification primers designed from conserved regions of *tRNA-Phe* and *tRNA-Ile* amplified templates suitable for sequencing from many different oomycete species (Martin et al., 2014; F. N. Martin, unpublished). While this locus was useful for phylogenetic analysis and as a barcode for species identification, a mean length of approximately 600 bp makes it too long for metabarcoding using Illumina sequencers. Yuan et al. (2017) also noted the sequence divergence of the *rps10* locus (among others) for 14 oomycete taxa and suggested the locus as a candidate barcode for oomycetes.

Here, we propose the *rps10* locus as an oomycete barcode and provide primers suitable for short-read sequencing on platforms like the Illumina MiSeq. The usefulness of *rps10* primers was compared to the semi-nested method to amplify ITS1 proposed by Sapkota and Nicolaisen (2015), which has reported some of the best results using ITS1 for oomycete metabarcoding. We compared each method’s taxonomic specificity and resolution by simulating PCR amplification with reference sequences and conducting metabarcoding of environmental samples and a mock community composed of a mixture of known species composition. We clustered reads into both ASVs and OTUs to determine which approach works best with the proposed method. We developed a companion website to host the *rps10* reference database and describe all protocols needed for researchers to immediately apply this validated method to oomycete metabarcoding.

## 2. MATERIALS AND METHODS

### 2.1 Primer design

Metabarcoding primers suitable for use with the Illumina MiSeq (San Diego, CA) were designed by searching sequence alignments of the region around the *rps10* locus. The sequences consisted of amplicons generated using primers in the flanking *tRNA-Phe* and *tRNA-Ile* loci (F. N. Martin et al., 2014) or extracted from assembled mitochondrial genomes (F. N. Martin, unpublished) and represented 16 genera and 92 species of oomycetes. Sequences were aligned with Clustal Omega (Sievers et al., 2011) and inspected in Geneious 8.1.9 (Biomatters, Auckland, New Zealand). Forward and reverse primers were designed for the highly conserved regions in the *tRNA-Phe* and *tRNA-Arg* genes flanking the *rps10* gene (Figure 1). Pairs of potential primers for metabarcoding were chosen that would: (1) result in an amplicon length of less than 500 bp and would therefore be appropriate for short-read sequencing using platforms like the Illumina MiSeq, (2) be conserved in all known oomycetes, and (3) not match the sequences of other organisms, particularly fungi and plants (Cristescu, 2014). Potential primers were evaluated with OligoAnalyzer (Owczarzy et al., 2008) to assess their melting temperatures, CG content, and potential for forming problematic secondary structures, such as dimers and hairpins. Table 1 provides the primers developed for *rps10* metabarcoding and the sequences found in the reference database.

**Figure 1.**
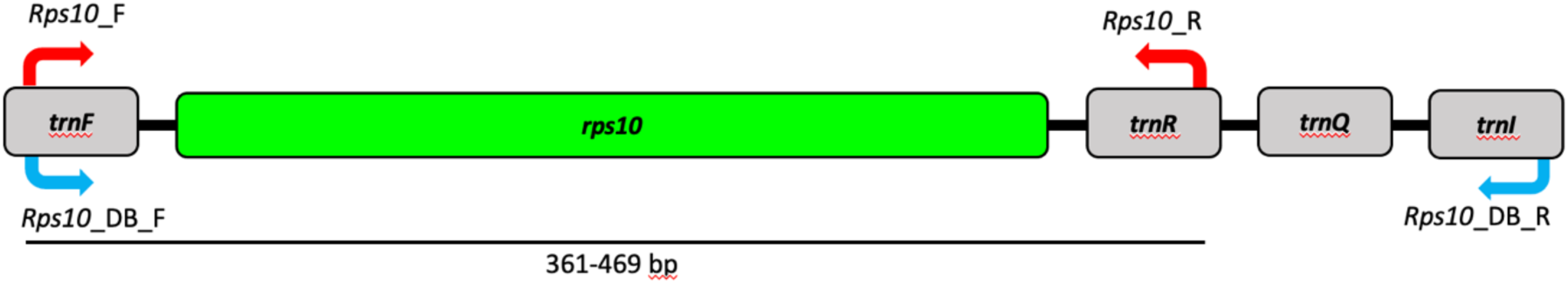
The location of the 40S ribosomal protein S10 (*rps10*) locus in the mitochondrial genome of oomycetes. The *rps10* locus is flanked by several tRNA genes (F. N. Martin, Bensasson, Tyler, & Boore, 2007). Primers shown in blue are for amplification of the sequences found in the reference database, and primers shown in red are for metabarcoding. More details on the two sets of primers are provided in Table 1.

**Table 1.**
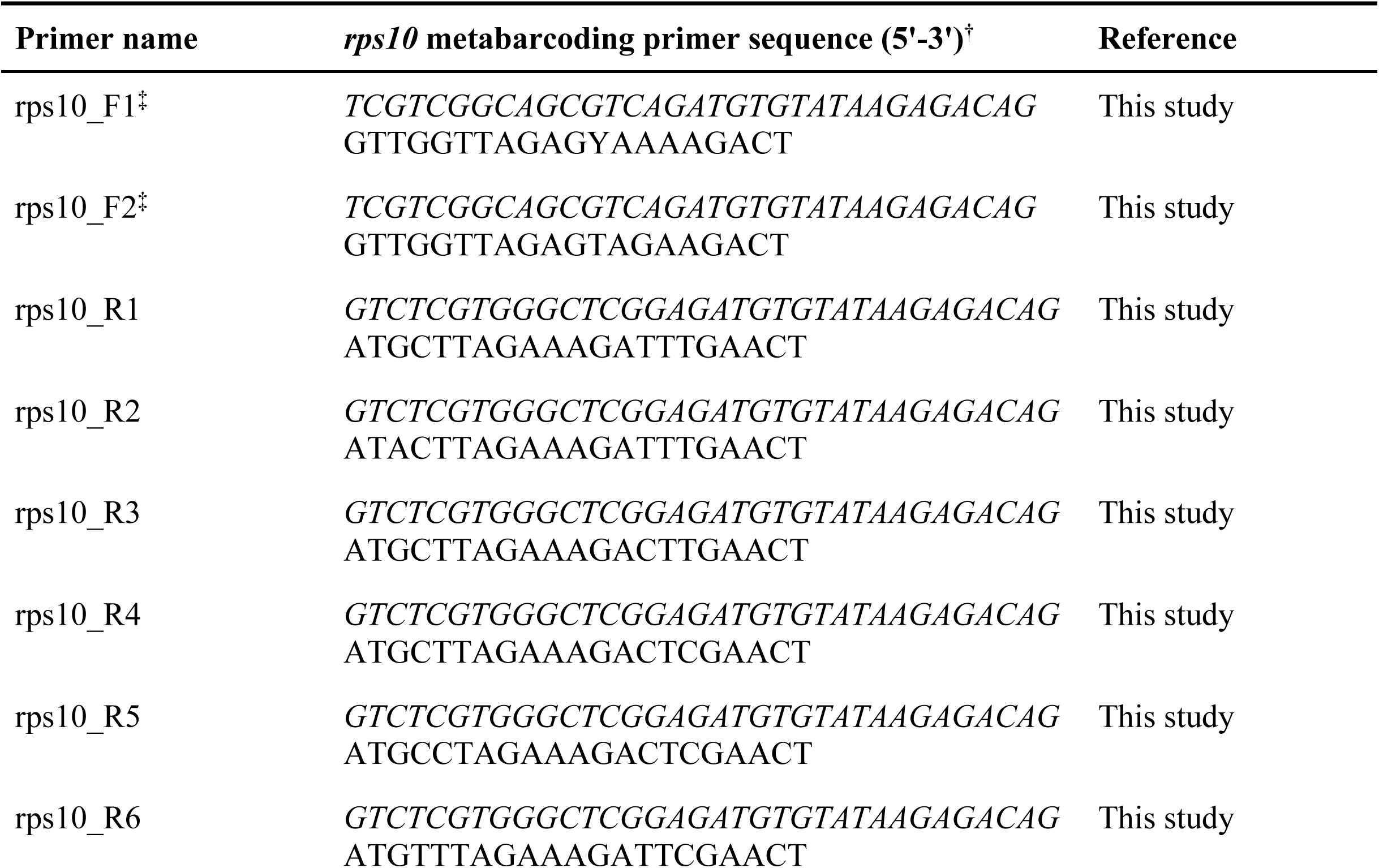
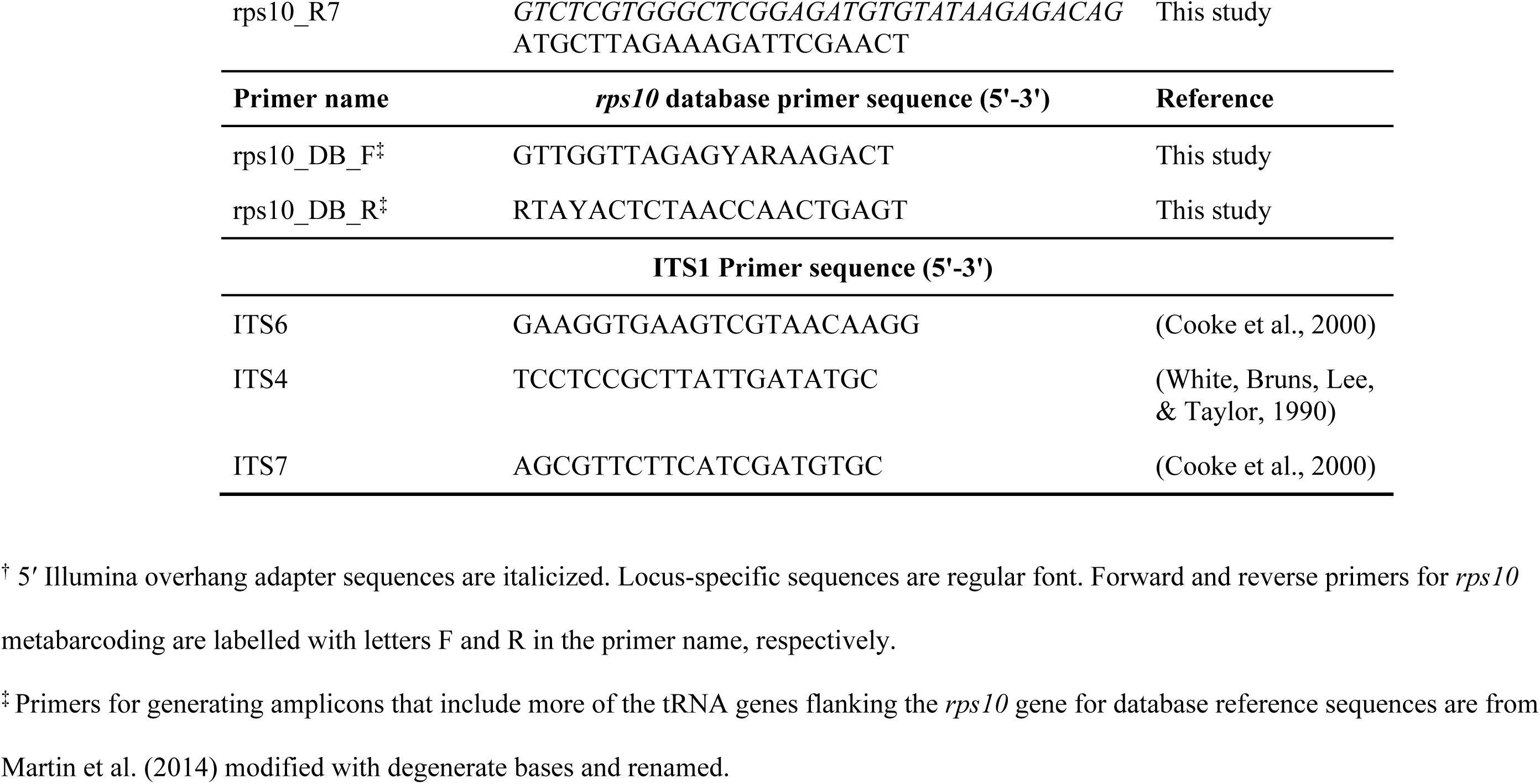
List of primers used for *rps10* metabarcoding and the reference database entries.

### 2.2 Simulated PCR

To test the newly developed primers for taxonomic specificity and coverage, we used the *rps10* locus extracted *in silico* from the mitochondrial genomes of 121 oomycetes, 38 non-oomycete stramenopiles, 19 fungi from different families, and four *Rickettsia* species. *Rickettsia* species were included because segments of their genomes resemble mitochondrial genomes (Andersson et al., 1998). We evaluated the sensitivity and specificity of the primers using Geneious 8.1.9 with the following parameters: no mismatches allowed, SantaLucia (1998) formula and salt correction, 50 nM concentration of oligos, and 0.6 mM dNTPs. The results were analyzed and visualized in a taxonomic context using the R packages *taxa* (Foster, Chamberlain, & Grünwald, 2018) and *metacoder* (Foster, Sharpton, & Grünwald, 2017).

### 2.3 Isolate and environmental DNA collection

To test for amplification of oomycetes and taxonomic resolution using the metabarcoding primers, a DNA mixture representing a mock community of 24 oomycete species was prepared (Table S1). The mock community was composed of a mixture of known oomycete species from different laboratories. The DNA was extracted from cultured strains or host tissue (for obligate pathogens) depending on species (Table S1). DNA from each of the 24 species was pooled, resulting in a per-species final concentration of ∼2 ng/µL, with the exception of a lower concentration for *Saprolegnia diclina* (0.5 ng/µL) and *Phytophthora pluvialis* (1.0 ng/µL). The concentration of DNA measured represented all the DNA in the sample; thus, DNA extracts from infected plant tissue (for obligate pathogens) included an unknown proportion of plant DNA and an unknown concentration of oomycete DNA.

To test for non-target amplification, we extracted DNA from diverse environmental samples including soil, water, and plant tissue. Samples consisted of soils from Panama (Schappe et al., 2017) and soil, canopy drip water, and tree needles collected near old-growth *Pseudotsuga menziesii* in the Wind River Forest Dynamics Plot in Washington State, USA. DNA from soil was extracted following a previously reported protocol (Schappe et al., 2017). DNA from needles was extracted using the DNeasy Plant Mini Kit (Qiagen, Valencia, CA, USA), following the manufacturer’s instructions. Water samples were collected by filtering up to 1L of water dripping through tree canopies with 0.45 µm filters and extracting DNA from the filters using the DNeasy Plant Mini Kit.

### 2.4 DNA amplification and high-throughput sequencing

DNA from the mock community and environmental samples was amplified with both the *rps10* method and the ITS1 method (Sapkota & Nicolaisen, 2015) to generate amplicons for high-throughput sequencing. Negative PCR water controls were included in both assays. The *rps10* assay is a multiplex PCR reaction comprising two *rps10* forward primers that differ slightly in sequence but anneal to the same position in the *tRNA-Phe* gene (rps10_F1 and rps10_F2) and seven *rps10* reverse primers that differ slightly in sequence but anneal to the same position in the *tRNA-Arg* gene (rps10_R1 through rps10_R7) (Table 1). All amplifications of the *rps10* locus were carried out using the QIAGEN Type-it Mutation Detect PCR Kit (QIAGEN, 206343, Valencia, CA). Multiplex PCR reactions were performed in 35 µL with 14 ng template DNA and 1 X final buffer concentration. Final primer concentrations were 0.2 µM except for rps10_F1, which has a single degenerate base (Y = C, T) representing two primers and was therefore added at a final concentration of 0.4 µM. Amplifications were carried out in a Veriti thermal cycler (Life Technologies, Grand Island, NY) with an initial denaturation at 95 °C for 5 min, followed by 35 cycles of 95 °C for 30 s, 58 °C for 3 min, and 72 °C for 30 s, and a final extension at 60 °C for 30 min.

ITS1 was amplified following a semi-nested protocol with minor modifications from the previously published method using the ITS6/ITS4 and ITS6/ITS7 primer sets (Sapkota & Nicolaisen, 2015). The initial PCR reaction was performed in 1X PCR, 1.5 mM MgCl_2_, 0.2 mM dNTP mix, 0.2 µM of each primer (ITS6 and ITS4), and 2 U/reaction of Platinum *Taq* (#10966018, ThermoFisher Scientific). The non-proofreading Platinum *Taq* was used since previous efforts to use various proofreading *Taq* polymerases resulted in unacceptably strong amplification of plant DNA using these primers (unpublished data), possibly due to the 3’ to 5’ exonuclease activity of proofreading *Taq* polymerase removing bases at the 3’ end of the ITS7 primer that distinguishes oomycetes from plants (Sapkota & Nicolaisen, 2015). DNA (30 ng, except for *P. pluvialis* at 15 ng and *S. diclina* at 7.5 ng) was added to each reaction for a total volume of 15 µL. The second PCR reaction was identical to the first with the following exceptions: the template was 1.0 µL of the initial PCR reaction, primers were ITS6 and ITS7, and the total reaction volume was 25 µL. Both ITS1 PCR amplifications were conducted in a Bio-Rad T100 thermocycler (Bio-Rad, Hercules, CA, USA) under the following thermal cycling conditions: 2 min at 94°C, 25 cycles of 30 sec at 94°C, 30 sec at 60°C, 1 min at 72°C, and a final extension of 2 min at 72°C. The protocol used differs slightly from that of Sapkota and Nicolaisen (2015); the first PCR had 25 cycles instead of 15 and the annealing temperatures of the PCR were raised from 55 °C and 59 °C to 60 °C for both. These changes were based on optimizing the PCR conditions to minimize non-target amplification in a previous experiment (Foster, Weiland, Scagel, & Grünwald, 2020). Amplicons from both the *rps10* and second ITS1 PCR amplicons were then cleaned, ligated to Illumina Nextera XT indices and adapters, purified, and pooled following Illumina manufacturer’s protocols (Illumina, 2013). Sequencing was carried out with 300-bp paired-end reads on the Illumina MiSeq platform at the Center for Genome Research and Biocomputing at Oregon State University.

### 2.5 *Rps10* database and associated website

A curated reference database was developed for assigning taxonomic classifications to sequences generated from the *rps10* primers. The database is composed of sequences manually curated from online databases, contributed by other research groups, identified from whole mitochondrial genomes, and produced from known isolates in this study. Species classification was confirmed by *cox1* or ITS sequencing and comparison with sequences from vouchered specimens in GenBank. To generate *rps10* sequences for the reference database we attempted to find primers that would produce an amplicon that included both metabarcoding primer binding sites, which would be useful for detecting primer mismatches when new organisms are sequenced in the future, but found it was only possible to include the binding site of the reverse primer. We selected previously reported amplification primers (F. N. Martin et al., 2014) and modified them by addition of degenerate bases. Amplification of the *rps10* database amplicon was conducted in 25.0 µL reactions with 0.025 U/µL GenScript Taq (GenScript, Cat. No. E00007), 1 X Taq Buffer, 0.2 µM dNTPs, 1.5 mM MgCl_2_, 2.0 ng DNA, and 0.5 µM of each primer (rps10_DB_F and rps10_DB_R). Thermal cycling was performed using a Veriti thermal cycler with an initial denaturation at 94 °C for 3 min, followed by 35 cycles of 94 °C for 30 sec, 55 °C for 45 sec, and 72 °C for 45 sec, and a final extension at 72 °C for 7 min. This is the suggested protocol for future researchers to use to contribute sequences to the reference database.

### 2.6 Abundance matrix preparation

An ASV abundance matrix with associated taxonomic annotations was created from MiSeq reads using *cutadapt* (M. Martin, 2011) and the R package *dada2* (Callahan et al., 2016). Primer sequences were trimmed from reads using *cutadapt*. Reads were then filtered out using the *filterAndTrim* command of *dada2* if they were expected to contain 5 or more errors, based on their quality scores. Reads were also truncated at the first instance of a quality score less than 5. Error rates were estimated and used to infer ASVs using the *learnErrors* and *dada* commands for each locus. ASV read pairs were merged using the *mergePairs* function and predicted chimeras were removed using *removeBimeraDenovo*. Any merged sequences less than 50 bp long were also removed. A taxonomic classification was assigned to each ASV using the RDP Naive Bayesian Classifier algorithm implemented in the *assignTaxonomy* command (Wang, Garrity, Tiedje, & Cole, 2007). The algorithm assigns a bootstrap value to each taxonomic rank for each classification, providing a confidence measure for which taxon in the reference database is most similar. For *rps10* sequences, the newly developed *rps10* database described herein was used as the reference database; for ITS1, a combination of UNITE (Kõljalg et al., 2005), Phytophthora-DB (Park et al., 2008), and sequences from Robideau et al. (2011) were used. Each ASV was also optimally aligned to the best-matching reference sequence to calculate a percent identity using the *pairwiseAlignment* function from the *biostrings* R package (Pages, Aboyoun, Gentleman, & DebRoy, 2017).

Reads were also clustered into OTUs using VSEARCH (Rognes, Flouri, Nichols, Quince, & Mahé, 2016) to create an OTU abundance matrix. To inform the choice of clustering threshold for each locus, ASVs present in the mock community were clustered at a range of thresholds from 90% to 100% in 0.1% increments and the number of resulting OTUs was recorded. Thresholds were chosen that best reproduced the number of species used in the mock community and conformed with previous experience using ITS1 as an oomycete barcode (Foster et al., 2020). *Rps10* sequences were clustered at a 96% threshold and ITS1 sequences were clustered at a 99% clustering threshold. Chimera detection, taxonomic assignment, and calculation of percent identity of OTUs to the assigned reference sequence were done with the same methods used for ASVs described above.

### 2.7 Mock community

The inferred composition of the mock community based on sequencing results was compared with the known composition of the mock community to evaluate the performance of the *rps10* and ITS1 methods. For this analysis, only ASVs represented by at least 10 reads were used. ASVs found in the mock community samples were classified as “expected”, or “non-target”. Three metrics were used to evaluate the ability of each method to infer the composition of the mock community: (1) the number of mock community members detected, (2) the proportion of ASVs representing members of the mock community, and (3) the proportion of reads representing members of the mock community. These three metrics were calculated using both the taxonomic classifications and the sequences, resulting in a total of 6 metrics for each locus. Taxonomic classifications were considered correct that contained the species name of a member of the mock community. Sequences were considered correct if they matched the reference sequences of the mock community members exactly, as determined by alignments of each ASV to each reference sequence using the *pairwiseAlignment* function from the *biostrings* R package. The same analysis was done allowing for a single mismatch in an alignment between the ASV and the reference sequence to check for nearly correct sequences.

### 2.8 Taxonomic specificity

ASVs and OTUs associated with environmental samples were used to assess non-target amplification. Since our reference databases do not contain sequences for many groups of potential non-target organisms, ASVs and OTUs in this analysis were assigned an alternative taxonomic classification using a BLAST search against the NCBI nucleotide database (Altschul, Gish, Miller, Myers, & Lipman, 1990). Although NCBI taxonomic annotations can be unreliable (Nilsson et al., 2006), we only considered the kingdom-level part of the taxonomy, which we considered more likely to be correct. The best BLAST hit was chosen for each ASV/OTU based on the E-value and percent identity of the matching region. BLAST hits with an E-value higher than 0.001 were not considered. Using the taxonomy associated with the best BLAST hit, ASVs were grouped into “Oomycetes”, “Fungi”, and “Other” categories. They were considered “Unknown” when no acceptable BLAST hit was found. The proportions of reads, ASVs, and OTUs in each category for each locus were then compared to evaluate the amount of non-target amplification for each locus.

### 2.9 Taxonomic resolution

The ability of each locus to distinguish different species was evaluated by pairwise alignments of reference database sequences. The portion of the reference database sequences predicted to be amplified by each primer pair was aligned using MAFFT v7.453 (Katoh, 2005) and all pairwise differences in sequence identity were calculated using the *ape* R package. Reference sequences with the complete amplicon were used, as determined by the presence of primer binding sites, using a modified version of the *matchProbePair* function from the *Biostrings* package that allows for ambiguity codes. Reference sequences were also included if they aligned to at least 90% of one of the predicted amplicons, in which case the portion aligned to the amplicon was used. Only reference sequences for species present in both databases were used in order to make the comparison as fair as possible. For each locus, the distributions of the percent identity of each species’ sequence to the most similar sequence from a different species were compared to assess taxonomic resolution.

The taxonomic resolution was also investigated by comparing the distribution of bootstrap scores of ASV taxonomic assignments and the bootstrap scores generated by the neighbor-joining tree of mock community species. The bootstrap values used were those assigned by the RDP Naive Bayesian Classifier algorithm implemented by the *assignTaxonomy* command of *dada2*; these values measure how consistent the taxonomic assignment is for a given reference database when parts of the sequences are subsampled. Bootstrap values were also generated to assess taxonomic resolution when there were multiple reference sequences for the same species. The distribution of bootstrap values for mock community samples for the genus, species, and reference sequence ranks were compared for ITS1 and *rps10*. In addition, the bootstrap scores for clades in the neighbor-joining trees of mock community species were compared.

## 3. RESULTS

### 3.1 Validation of metabarcoding primers for the *rps10* region

The new primers developed for oomycete-specific amplification of the *rps10* locus were validated using simulated PCR of reference sequences (Figure 1). There are two forward primers and seven reverse primers that bind to the same respective regions but differ slightly in sequence (Table 1). Simulated PCR using the forward and reverse primer sets amplified the *rps10* gene with no mismatches on any of the 121 oomycete sequences analyzed (Figure 2). In addition, the *rps10* primers were predicted not to amplify any non-target species found in the stramenopiles, fungi, and *Rickettsia* (Figure 2). The forward primer set has 40% mean GC content and a predicted mean melting temperature of 59.8°C, while the reverse primer set has 31% mean GC content and a predicted mean melting temperature of 58.2°C. The primers are predicted to amplify a sequence with a median length of 481 bp (including primers), with *Albugo laibachii* producing the shortest with 448 bp, and *Peronospora tabacina* producing the longest with 513 bp (Figure S1). The length of amplicons produced by the ITS1 method, inferred using the same procedure, varied from 225 bp to 400 bp.

**Figure 2.**
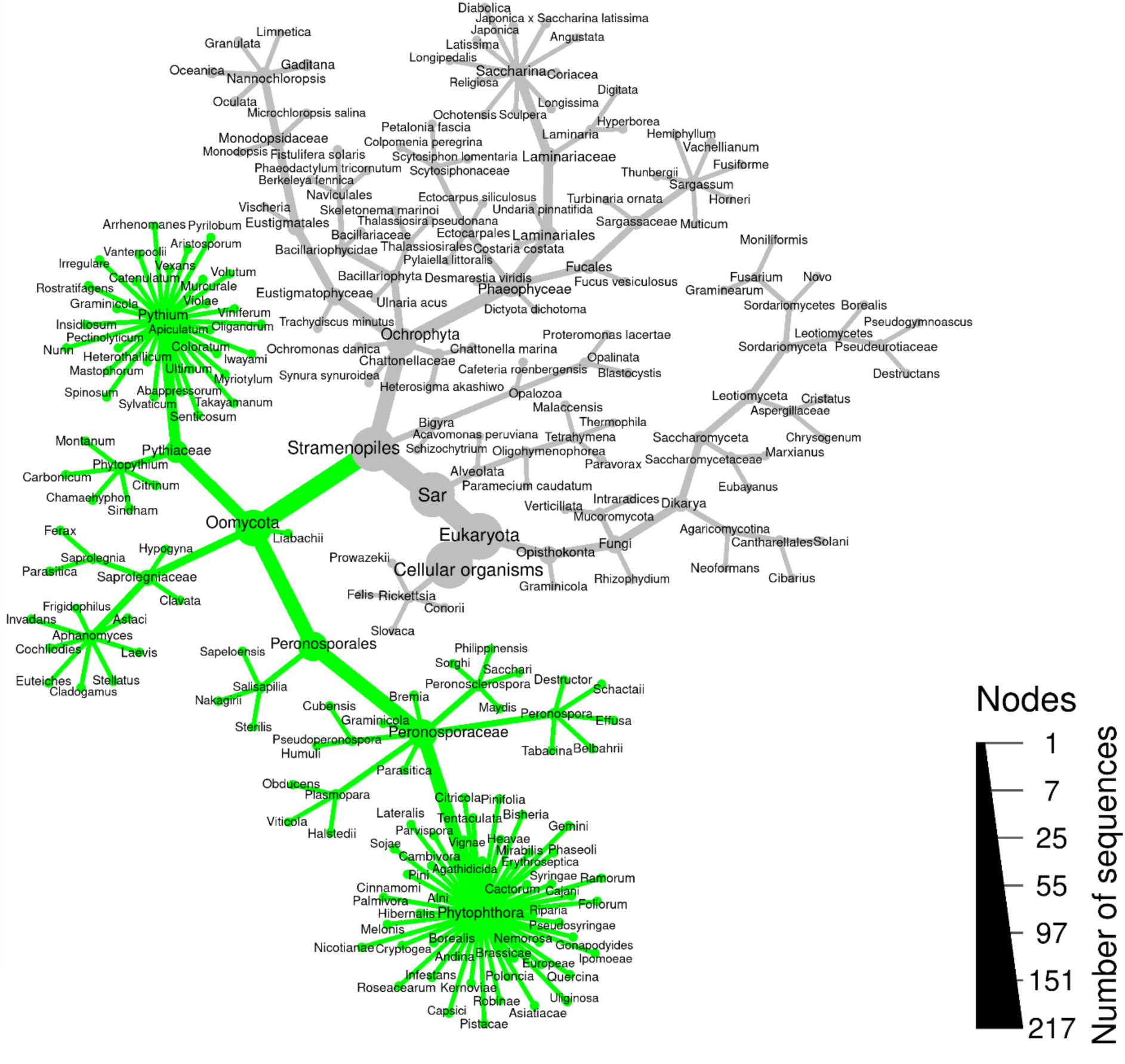
Heat tree showing predicted amplification (green) of the *rps10* metabarcoding primers. Taxa in green are predicted to be amplified using simulated PCR. The analysis included fungi and related Stramenopiles to demonstrate specificity. *Rickettsia* were also included since their genomes resemble mitochondrial genomes. The size of branches and nodes is relative to the number of sequences represented by each taxon.

### 3.2 The *rps10* reference database and website

To host the *rps10* database and laboratory protocols, we created the website www.oomycetedb.org. The database currently contains 886 sequences representing 346 species and 20 genera of oomycetes (Table 2). The website is a combination of static HTML produced with R markdown (Xie, Allaire, & Grolemund, 2019) and R shiny applications (www.rstudio.com/shiny/). Laboratory protocols can be viewed on the website or downloaded as printer-friendly PDFs. Users can download all or a specific subset of the database based on a search term. Users can also conduct BLAST (Altschul et al., 1990) searches of the database with their own sequences, view the results online, and download the results in any format BLAST can output. Updates to the database are released on this website with a unique version number and old versions will continue to be available for the sake of reproducibility. All tools on the website can be used with any version of the database so researchers can reproduce analyses. All source code and documents for the website are available on Github at <u>github.com/grunwaldlab/OomyceteDB</u>.

**Table 2.** Overview of the number of sequences currently available in the *rps10* barcode database.

| **Genus** | **Number of sequences** |
| --- | --- |
| *Achlya* | 1 |
| *Albugo* | 1 |
| *Aphanomyces* | 13 |
| *Bremia* | 3 |
| *Halophytophthora* | 9 |
| *Hyaloperonospora* | 1 |
| *Lagena* | 1 |
| *Lagenidium* | 2 |
| *Perofascia* | 1 |
| *Peronosclerospora* | 4 |
| *Peronospora* | 73 |
| *Phytophthora* | 509 |
| *Phytopythium* | 25 |
| *Plasmopara* | 5 |
| *Pseudoperonospora* | 3 |
| *Pythium* | 226 |
| *Salisapilia* | 4 |
| *Saprolegnia* | 3 |
| *Sclerospora* | 1 |
| *Thraustotheca* | 1 |
| **Total** | **886** |

### 3.3 Metabarcoding of the mock community

A mock community composed of DNA from 24 species of known oomycetes was sequenced on the Illumina MiSeq to evaluate the ability of the proposed *rps10* metabarcoding method to infer community composition relative to an ITS1-based method. Both the *rps10* and ITS1 methods resulted in more ASVs and OTUs than species included in the mock community. The *rps10* method resulted in fewer ASVs than the ITS1 method but more OTUs, suggesting that the ASV-based analysis better controlled for sequencing error than the OTU-based analysis (Table 3). The taxonomic classifications of ASVs produced by the *rps10* method included 23 of the 24 mock community species whereas the ITS1 classifications included 17 (Table 3). The species missing in the *rps10* classifications was *Phytophthora ipomoeae* and the species missing in the ITS1 classifications were *Phytophthora citrophthora*, *Phytophthora himalsilva*, *Phytophthora ipomoeae*, *Phytophthora quercina*, *Pythium dissotocum*, *Pythium oligandrum*, and *Pythium undulatum*. The *rps10* ASVs assigned to species in the mock community accounted for 95% of all ASVs and >99.9% of the reads, whereas the ITS1 ASVs with correct classifications accounted for 75.6% of all ASVs and 54.5% of the reads. Similar metrics were produced by considering whether the correct sequences were found, regardless of taxonomic classification. Both methods resulted in ASV sequences matching 21 of the 24 species exactly and matching all 24 when a 1 bp mismatch in the alignment between ASVs and reference sequences was tolerated. The *rps10* ASVs with exact sequence matches accounted for 62.5% of ASVs and 99.3% of reads, whereas the ITS1 ASVs with correct sequences matches accounted for 73.3% of ASVs and 99.2% of reads. In five out of the six metrics used to evaluate how well the mock community composition was inferred, the *rps10* method performed equal to or better than the ITS1 method (Table 3).

**Table 3.**
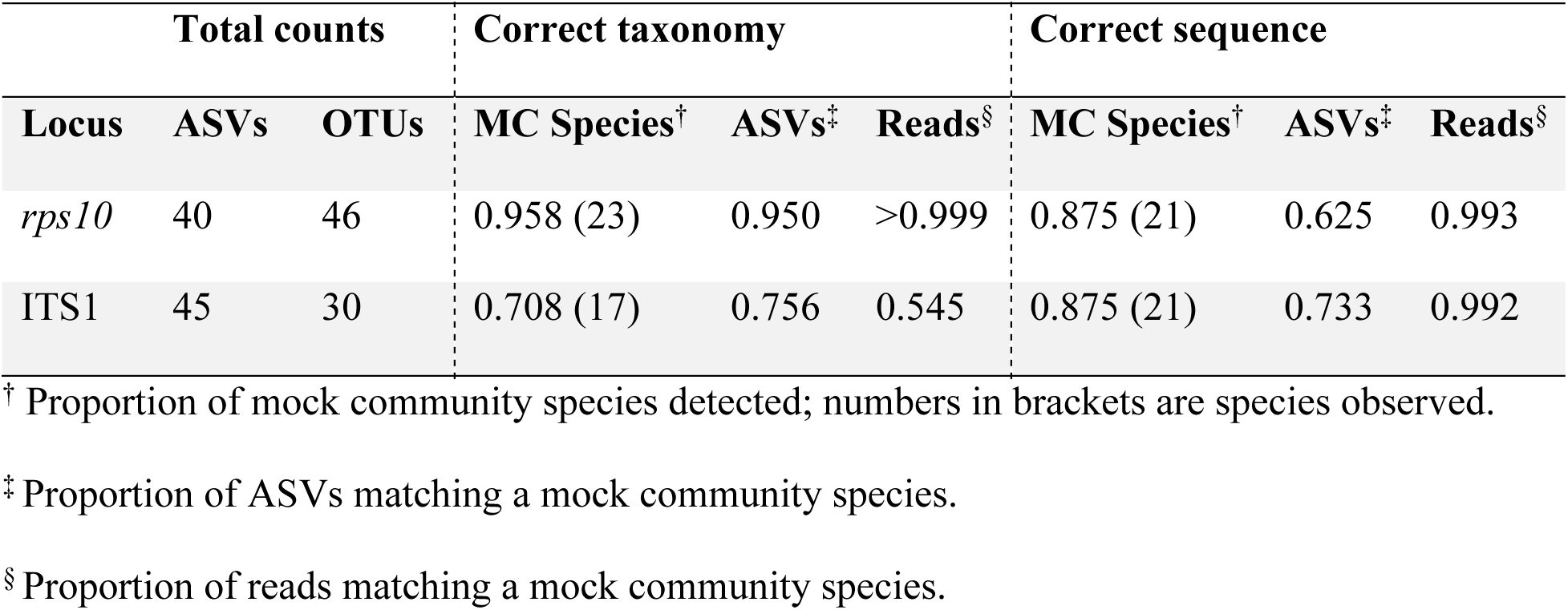
Mock community inference statistics based on proportions of correct taxonomic classifications and proportions of correct sequences.

| Total counts |  |  | Correct taxonomy |  |  | Correct sequence |  |  |
| --- | --- | --- | --- | --- | --- | --- | --- | --- |
| Locus | ASVs | OTUs | MC Species<sup>†</sup> | ASVs<sup>‡</sup> | Reads<sup>§</sup> | MC Species<sup>†</sup> | ASVs<sup>‡</sup> | Reads<sup>§</sup> |
| *rps10* | 40 | 46 | 0.958 (23) | 0.950 | >0.999 | 0.875 (21) | 0.625 | 0.993 |
| ITS1 | 45 | 30 | 0.708 (17) | 0.756 | 0.545 | 0.875 (21) | 0.733 | 0.992 |

<sup>†</sup> Proportion of mock community species detected; numbers in brackets are species observed.
<sup>‡</sup> Proportion of ASVs matching a mock community species.
<sup>§</sup> Proportion of reads matching a mock community species.

### 3.4 Metabarcoding of environmental samples

A diverse set of environmental DNA extracts were sequenced on the Illumina MiSeq using the proposed *rps10* metabarcoding method to evaluate taxonomic specificity. For the *rps10* method, oomycete sequences accounted for 99.5% of the reads, 84.8% of the ASVs, and 53.5% of the OTUs (Figure 3). For the ITS1 method, oomycete sequences accounted for 63.0% of the reads, 10.7% of the ASVs, and 5.5% of the OTUs. Most of the *rps10* sequences not assigned to oomycetes could not be assigned to a taxon based on BLAST searches against the NCBI nucleotide database, particularly for the plant-derived samples. Most of the ITS1 sequences not assigned to oomycetes were assigned to fungi and a much smaller proportion were assigned to plants. For both methods non-target ASVs and OTUs were at lower abundance than oomycete sequences, but this trend was more pronounced for *rps10*. Manual investigation of the *rps10* ASVs with no matches to NCBI revealed that these sequences are much shorter than typical *rps10* sequences, have a much higher GC content, and do not align well to *rps10* reference sequences or to each other. Overall, sequencing of amplicons from environmental samples from water, soil, and plant material using the *rps10* barcode resulted in less non-target amplification, in terms of ASV, OTU, and read counts than when using the ITS1 barcode (Figure 3).

**Figure 3.**
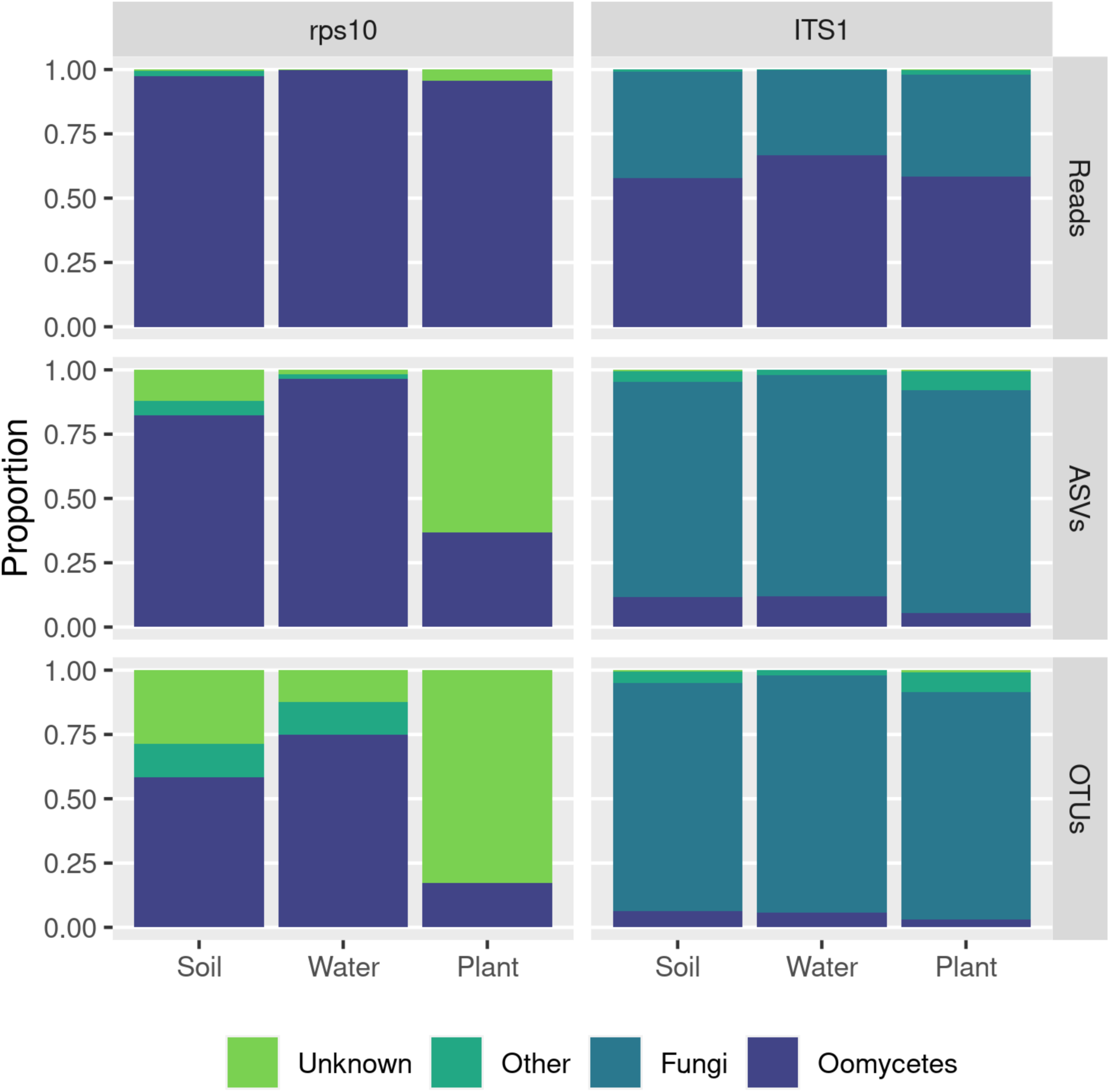
Taxonomic specificity of amplification using the *rps10* and ITS1 methods based on environmental samples. Counts of read, amplified sequence variants (ASVs), and operational taxonomic units (OTUs) from a variety of environmental samples grouped into soil, water, and plant tissue samples are shown. Here, ASV sequences were given a taxonomic assignment based on BLAST searches against the NCBI nucleotide database.

### 3.5 Taxonomic resolution

The ability to differentiate closely related species was evaluated by comparing how many base pairs differentiated each reference sequence from the most similar sequence for a different species in the region amplified. In general, the pairwise differences between the most similar sequences from different species were greater for *rps10* than ITS1 (Figure 4A). A total of 14.7% of the predicted amplicons derived from unique species in the *rps10* reference database shared an identical sequence with a different species, compared with 29.4% in the ITS1 database (Figure 4B). In addition, 67.6% of the species in the *rps10* database were distinguished from their most closely related species by five or more base pairs, in contrast to only 16.2% for ITS1. Polymorphic sites in the *rps10* locus are distributed across the entire length of the amplicons and sequences align with few gaps. In contrast, variation in the ITS1 locus appears in clusters and alignments have frequent large indels in many blocks (Figure 5). Another way to compare the ability to distinguish sequences is the distribution of bootstrap scores for taxonomic assignment and sequence clustering of the mock community sequences. There was little difference in the bootstrap scores for taxonomic assignment between the two methods. Since these bootstrap values are also influenced by the reference database and the reference databases were different for each locus, this comparison should be interpreted with caution. However, the bootstrap scores and branch lengths from the neighbor-joining trees of the mock community sequences were higher in *rps10* than in ITS1 (Figure S2; Figure S3), suggesting *rps10* sequences from different taxa are generally more diverged from each other than in similar comparisons for ITS1. Overall, *rps10* reference sequences exhibited more polymorphic sites among the most similar inter-species pairwise comparisons than ITS1 sequences and were overall more polymorphic.

**Figure 4.**
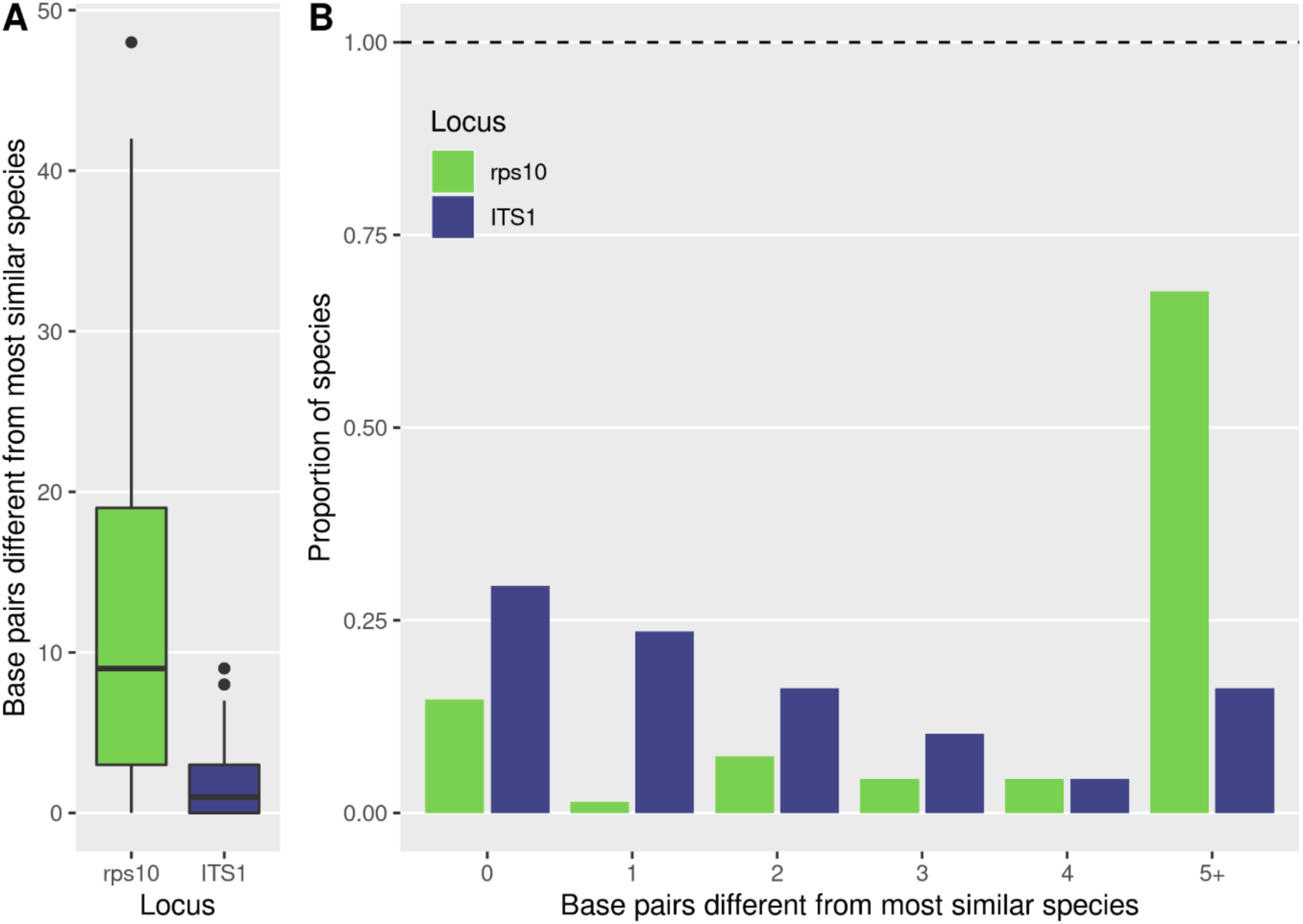
Taxonomic resolution of the *rps10* (green) and ITS1 (blue) barcodes. We evaluated the number of base pair differences to the most similar species based on pairwise alignments of predicted amplicons. (A) Distribution of the number of base pair differences between the most similar species. (B) Number of polymorphic sites differentiating the most similar species. Zero differences for a species mean there is at least one other species predicted to have an identical amplicon sequence. Only sequences for species present in both databases are included.

**Figure 5.**
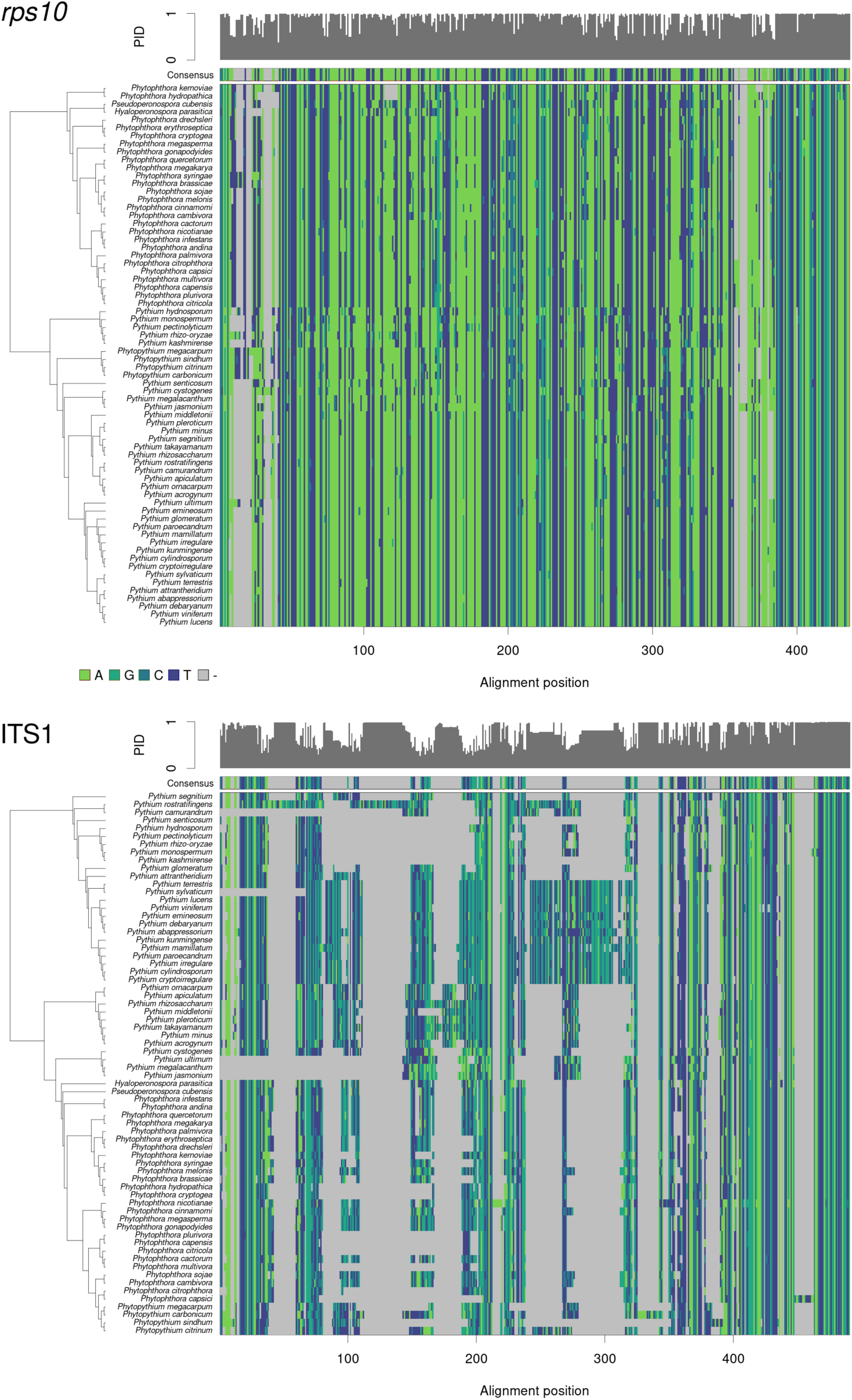
Multiple sequence alignments of the region predicted to be amplified by the *rps10* or ITS1 method, not including the primer binding sites. The sequences used represent the subset of species present in both reference databases. The sequences are ordered vertically based on a neighbor-joining tree. Along the top of each alignment is a bar chart representing the proportion of sequences matching the consensus sequence at each alignment position. Gray blocks are gaps in the alignment.

## 4. DISCUSSION

Effective methods for studying microbial communities without the need for culturing and manual identification have greatly increased our understanding of bacterial and fungal ecology, but methods for studying oomycetes are still in need of further development. Metabarcoding is one such method, but it requires a locus and associated primers with particular characteristics, including species-level taxonomic resolution, minimal length variation, the ability to amplify all target organisms, and minimal non-target amplification. The ITS1 locus is a popular choice for oomycete metabarcoding but suffers from limited taxonomic resolution and large variations in length. In addition, currently used primers targeting oomycetes often amplify non-target organisms as well. Here, we propose the *rps10* locus as a suitable metabarcoding locus for oomycetes and provide primers that produce an amplicon suitable for use with the Illumina MiSeq. Overall, our results suggest the *rps10* metabarcoding method results in less non-target amplification and has better taxonomic resolution than the ITS1 method tested and is best suited for the preferred ASV-based rather than OTU-based analysis.

### Taxonomic specificity

Simulated PCR of *rps10* reference sequences showed that all 121 oomycete sequences tested, representing 16 genera, should be amplified by the proposed primers and that non-target taxa, including non-oomycete stramenopiles, should not be amplified (Figure 2). Taxa amplified include species of global and economic concern belonging to the Saprolegniaceae and Peronosporaceae, such as *Aphanomyces euteiches*, *Phytophthora cinnamomi*, and *Phytophthora infestans*. We did not compare the *rps10* and ITS1 methods using simulated PCR because ITS1 sequences that included both primer binding sites for many of the species tested in this analysis are not publicly available. This is probably because many ITS1 reference sequences are produced with at least one of the primers we used (or primers binding to the same region) and therefore do not include the primer binding sites (Bellemain et al., 2010). However, our analysis of MiSeq data from diverse environmental samples suggested that the *rps10* method produced substantially less non-target amplification than the ITS1 method in terms of the proportion of non-target reads, ASVs, and OTUs (Figure 3). The unique order of tRNAs flanking the *rps10* gene is likely responsible for this level of amplification specificity (Figure 1). The non-target sequences from ITS1 were nearly all assigned to fungi (84.8% of ASVs, 36.3% of reads), which conforms to the results of other studies (Coince et al., 2013). Although Sapkota and Nicolaisen (2015) suggests increasing annealing temperature as a solution, we still observed much non-target amplification using the ITS1 method at several annealing temperatures (data not shown). Most of the non-target sequences from *rps10* had no close match to any sequence in the NCBI nucleotide database (12.1% of ASVs, 0.2% of reads). These unknown sequences tended to be low abundance, short, and highly dissimilar to *rps10* reference sequences, suggesting they may be erroneous. The minimal non-target amplification of the *rps10* method should result in more efficient use of sequencing throughput, making it possible to sequence communities in which oomycetes are rare relative to other organisms like fungi.

Sequencing of a mock community using both methods suggest they perform equally well at generating the correct sequences, but that the *rps10* method is better able to generate correct taxonomic classifications. Both methods produced amplicons that perfectly matched 21 of the 24 mock community species and all 24 when a single base pair mismatch was tolerated (Table 3). The ITS1 method produced a slightly greater proportion of ASVs matching mock community members (73.3% vs. 62.5%), but for both methods correct ASVs accounted for >99% of the reads. In terms of taxonomic classifications of ASVs, only 1 species out of 24 was not detected using the *rps10* method, in contrast to 7 missing with the ITS1 method. Considering that both methods produced sequences matching the same number of mock community members, the higher misclassification rate of the ITS1 method is likely due to insufficient taxonomic resolution. For example, both methods are missing *Phytophthora ipomoeae* from their taxonomic classifications but do include *Phytophthora infestans*. Since the amplified sequence for the two species is identical, it is likely that the amplicons for both species were assigned to *Phytophthora infestans*. This result suggests that the *rps10* region will produce more accurate taxonomic classifications when applied to real communities of unknown organisms even though the primers for the two methods amplify oomycetes equally well.

### Taxonomic resolution

Comparisons of the region predicted to be amplified in reference sequences suggest the *rps10* locus has greater taxonomic resolution than ITS1. Pairwise alignments of the predicted amplicons from the reference databases indicated 14.7% of oomycete species tested from the *rps10* database shared the exact sequence with at least one other species, compared with 29.4% of species from the ITS1 database (Figure 4). Since all sequencing methods have some degree of error, a single base pair difference in the targeted locus might be insufficient to distinguish species. If a difference of two base pairs is required to distinguish species reliably, the proportion of species not uniquely identified by *rps10* increased to 16.2%, while that of ITS1 increases to 52.9%. This suggests that most species can be confidently assigned to a species with *rps10*, even in the presence of sequencing errors. These results are corroborated by the greater branch lengths observed in neighbor-joining trees of mock community sequences (Figure S3), suggesting the average difference between sequences from different species is greater in *rps10*. Although *rps10* cannot distinguish all oomycetes species, it has a higher taxonomic resolution than the currently used ITS1 and does not contain the frequent large indels characteristic of ITS1, making it easier to create the multiple sequence alignments needed for many phylogenetic analyses (Figure 5). The superior taxonomic resolution of *rps10* will increase the confidence of species classifications, which will be especially helpful in cases where closely related species have different pathological or ecological implications.

### Practical considerations

The proposed *rps10* method has many attributes that should make it more effective than the ITS1 method tested, but also a few potential problems to consider. A single, rather than a nested, PCR step should reduce the chance of contamination, sequencing error, and lower the cost of reagents. The *rps10* amplicon also has much less variation in length (Figure S1), which should reduce the amount of read count bias (Nichols et al., 2018). However, we observed a higher rate of chimera formation in some *rps10* samples relative to ITS1, as well as more low-abundance, potentially erroneous sequences. This problem was addressed by filtering out chimeras and using an ASV-based analysis with the *dada2* package instead of an OTU-based analysis. For this reason and other known advantages (Callahan et al., 2017), we recommend using an ASV-based analysis when using the *rps10* method proposed here. Although the *rps10* amplicon for some oomycetes is near the upper limit for MiSeq 300bp paired-end sequencing, we were able to amplify and merge the paired-end reads of *Plasmopara obducens*, the longest amplicon in the mock community and the 5th longest in the reference database. Less amplification of non-target organisms should reduce the need to optimize PCR conditions as well as improve amplification efficiency of target organisms due to reduced competition for primers. The greater number of oomycete sequences relative to non-target sequences should allow for more samples for a given sequencing depth. This is a substantial improvement compared to the ITS1 method, in which specificity is very sensitive to the chosen annealing temperature of the PCR (Sapkota & Nicolaisen, 2015) and even after optimization suffers from non-target amplification of fungi. The greater taxonomic resolution of the *rps10* locus should result in a greater proportion of correct taxonomic classifications, which is desirable for monitoring pathogen occurrence. All resources needed to use the *rps10* method for metabarcoding oomycete communities using the Illumina MiSeq, including laboratory protocols and a reference database for taxonomic classification of results, are provided at www.oomycetedb.org.

### Future research

The global diversity of oomycetes is still largely unknown, with little knowledge of where invasive species come from or their native habitat ranges. This is underlined by results from our environmental samples, where many of the ASVs found had less than 90% similarity to a reference sequence, meaning ASVs could only be classified at the genus or family level. This is partially because the *rps10* reference database is incomplete and because many, if not most, oomycetes occurring in natural ecosystems have not yet been described. However, there are thousands of oomycete specimens in herbaria around the world that could be leveraged to improve this and other databases. A collaborative effort to sequence *rps10* barcodes from a wider range of oomycetes is in progress and will improve the accuracy and usefulness of this barcode. We encourage the oomycete community to submit *rps10* sequences for inclusion at www.oomycetedb.org, along with *cox1* sequences to confirm species classifications.

Oomycetes are an important but relatively understudied group of organisms. Understanding their diversity and distribution will help understand future outbreaks of destructive pathogens like *Phytophthora ramorum* and *Phytophthora infestans* and characterize the mostly unknown ecological niches of oomycetes in natural ecosystems. Assuming the species tested here are representative of oomycete diversity, many of the taxonomic classifications of oomycete microbiomes using ITS1 metabarcoding could be underestimating their true diversity and misclassifying closely related species. Much more research is needed to characterize the natural diversity of oomycetes. Currently, metabarcoding using Illumina sequencing is the most cost-effective technique (Tedersoo et al., 2019). We hope the method presented here will facilitate new insights into oomycete biodiversity and ecology, just as robust methods for metabarcoding of fungi and bacteria have revolutionized our understanding of those organisms in recent decades.

## ACKNOWLEDGEMENTS

We thank Steve Klosterman, Brett Smith, and Annette Dodge for providing reference DNA. The work was supported in part by: National Science Foundation grant # DEB-1542681 to NJG, FAJ, and BMT; the USDA-ARS CRIS Project 2072-22000-039-00D to NJG; Project 2038-22000-015-00D to FNM; and the USDA-ARS Floriculture Nursery Initiative research programs to NJG.

## AUTHOR CONTRIBUTIONS

NJG and FNM conceived and designed the study. FNM identified *rps10* as a candidate locus for metabarcoding. ZSLF, VJF, MML, FEA, FAJ, and NJG conducted field sampling. VJF, MML, ZSLF, and FEA validated the *rps10* locus. ZSLF analyzed sequencing data from metabarcoding. FNM, HDTN, TIB, CR, and NJG provided reference DNA or sequences for the reference database. ZSLF, VJF, MML, and NJG developed and maintain the oomycete-DB database and website. BMT, FAJ, and NJG were responsible for overall project design and obtained funding. All authors edited and approved the final version of the manuscript.

## DATA ACCESSIBILITY

All *rps10* sequences were submitted to GenBank (MZ365320-MZ365440) and metabarcoding data were submitted to the SRA (Bioproject PRJNA699663; Accessions SRR13658632-SRR13658679). A curated *rps10* database for taxonomic classification is publicly available at www.oomyceteDB.org for downloading. We encourage submission of new species to the database to improve this resource for the community (see instructions online). Reproducible code for the analysis presented here is available at https://github.com/grunwaldlab/rps10_barcode.

## Supporting Information

**Figure S1.**
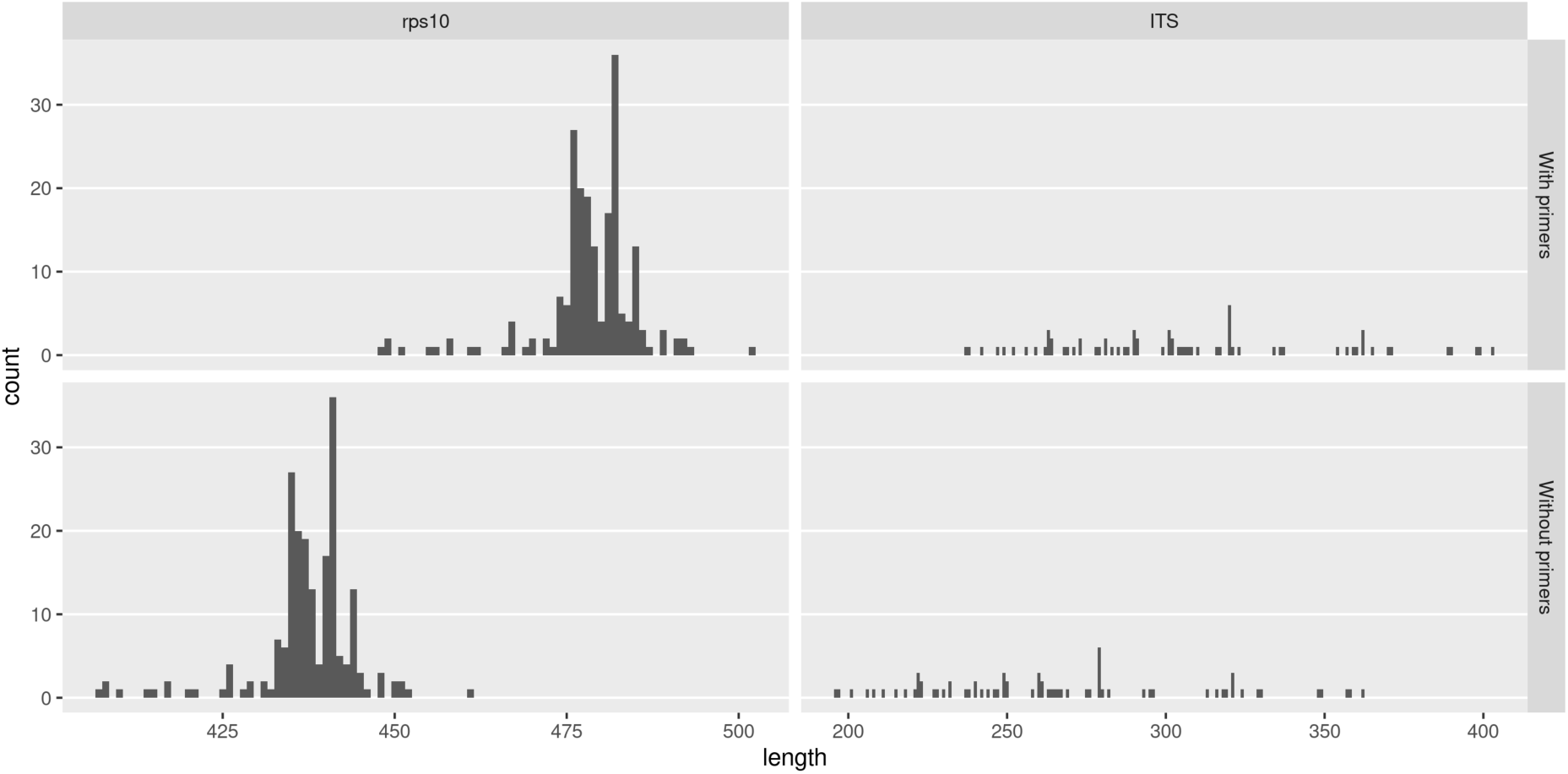
The distribution of amplicon lengths predicted to be produced by the *rps10* and ITS1 methods tested, with and without the primer sites (without the Illumina adapters), based on reference database sequences. Only species present in both databases are included.

**Figure S2.**
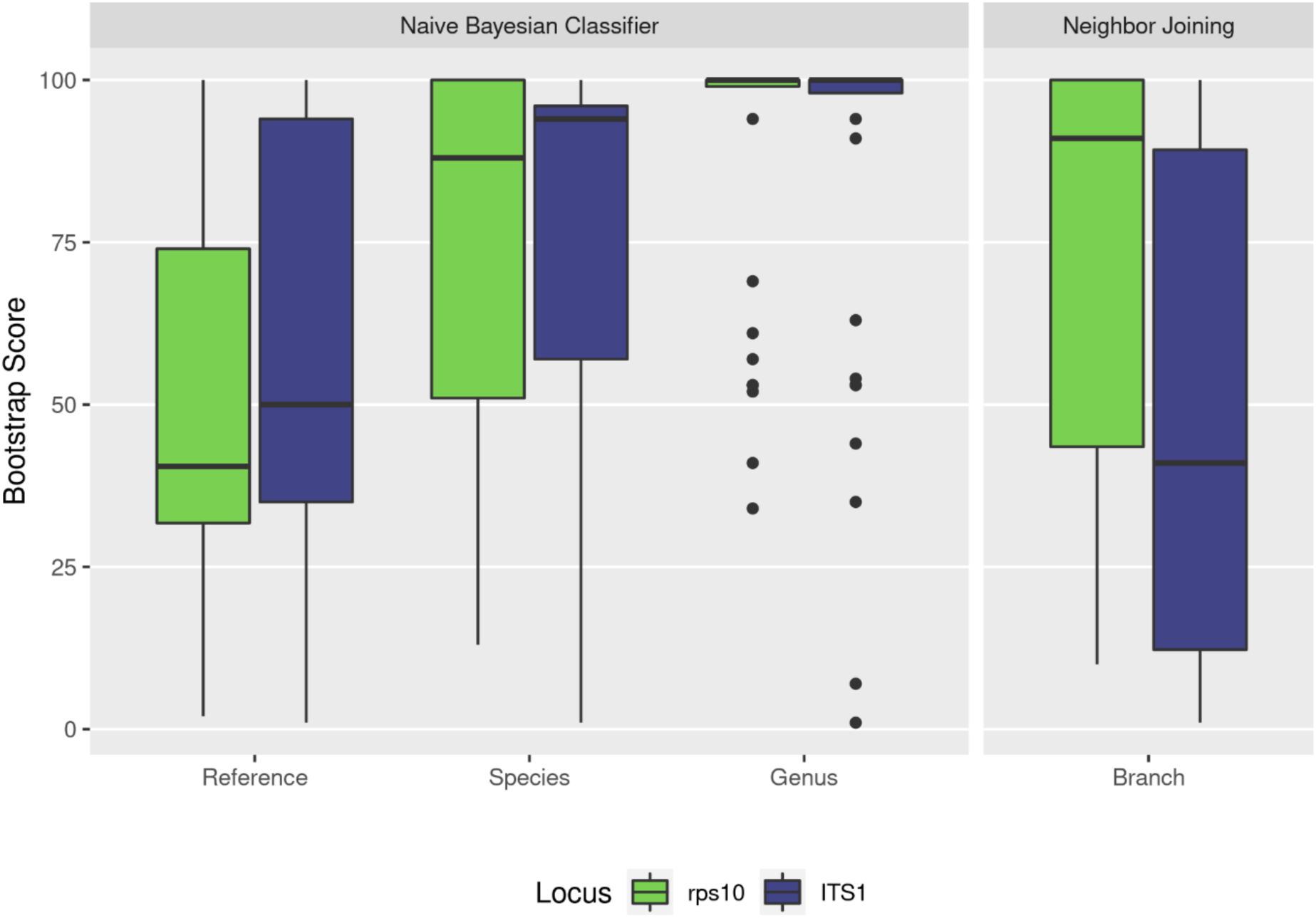
The distribution of bootstrap scores for the taxonomic assignment of amplified sequence variants (ASVs) in the mock community for the *rps10* and ITS1 loci. The RDP Naive Bayesian Classifier “Reference”, “Species”, and “Genus” scores refer to the ability to consistently assign ASVs to a particular reference sequence, species, or genus respectively when the data is resampled. The neighbor joining tree scores quantify how consistent the branching pattern of the resulting tree is when the data is resampled.

**Figure S3.**
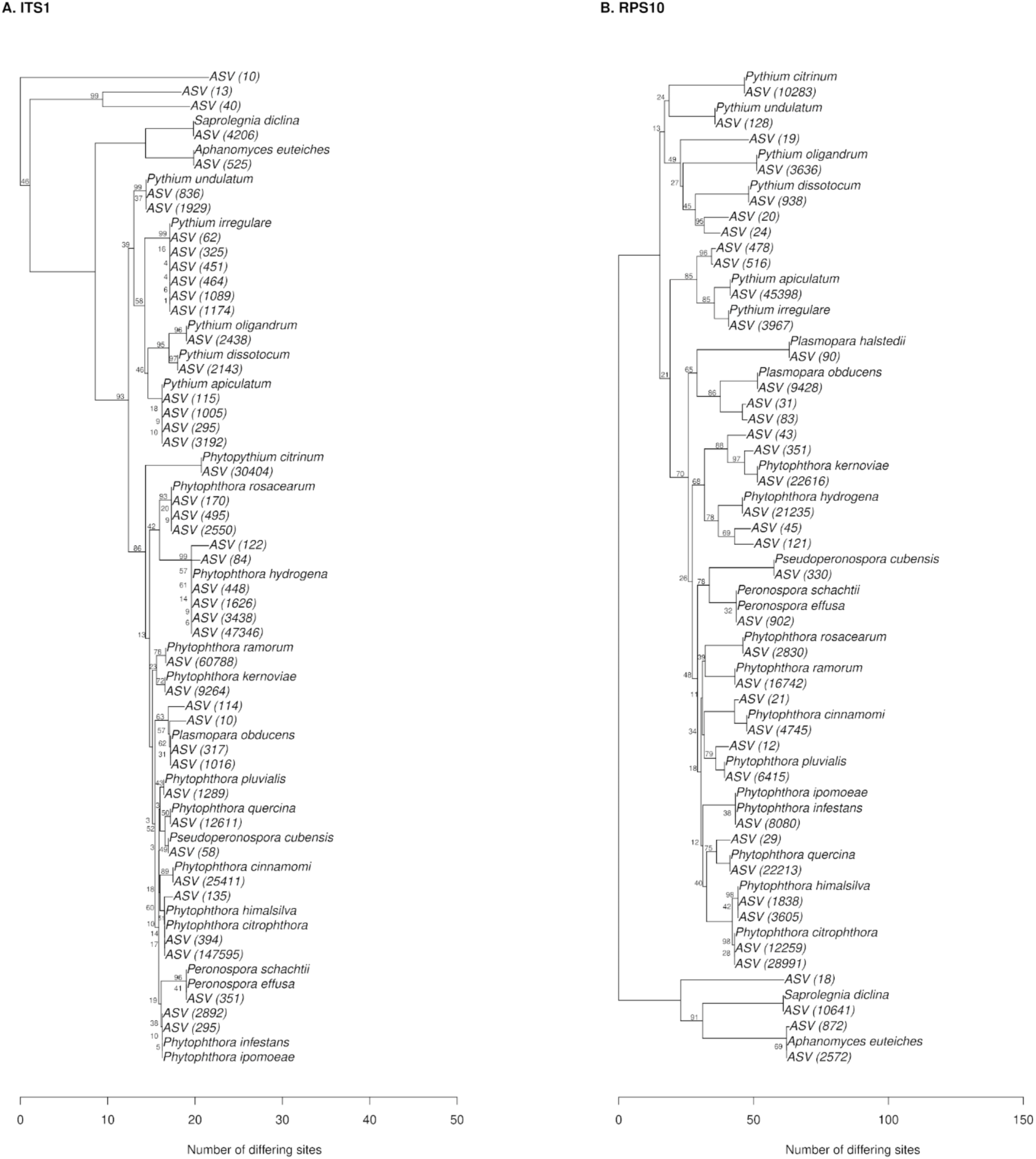
Bootstrapped neighbor-joining tree of ASVs in mock community samples and reference sequences for species included in the mock communities. The number of reads represented by each ASV is shown in parentheses.

---

**Table S1.**
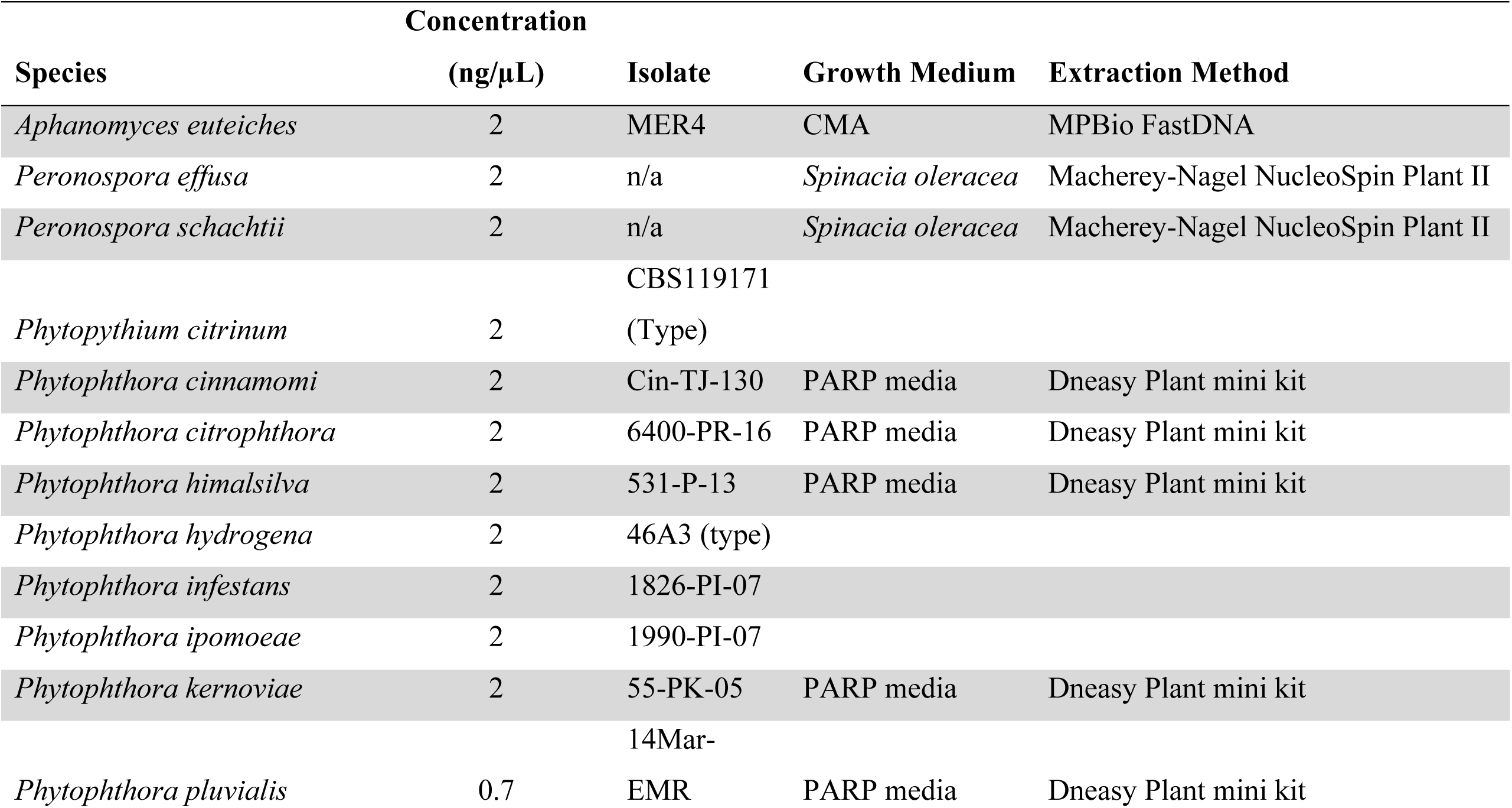
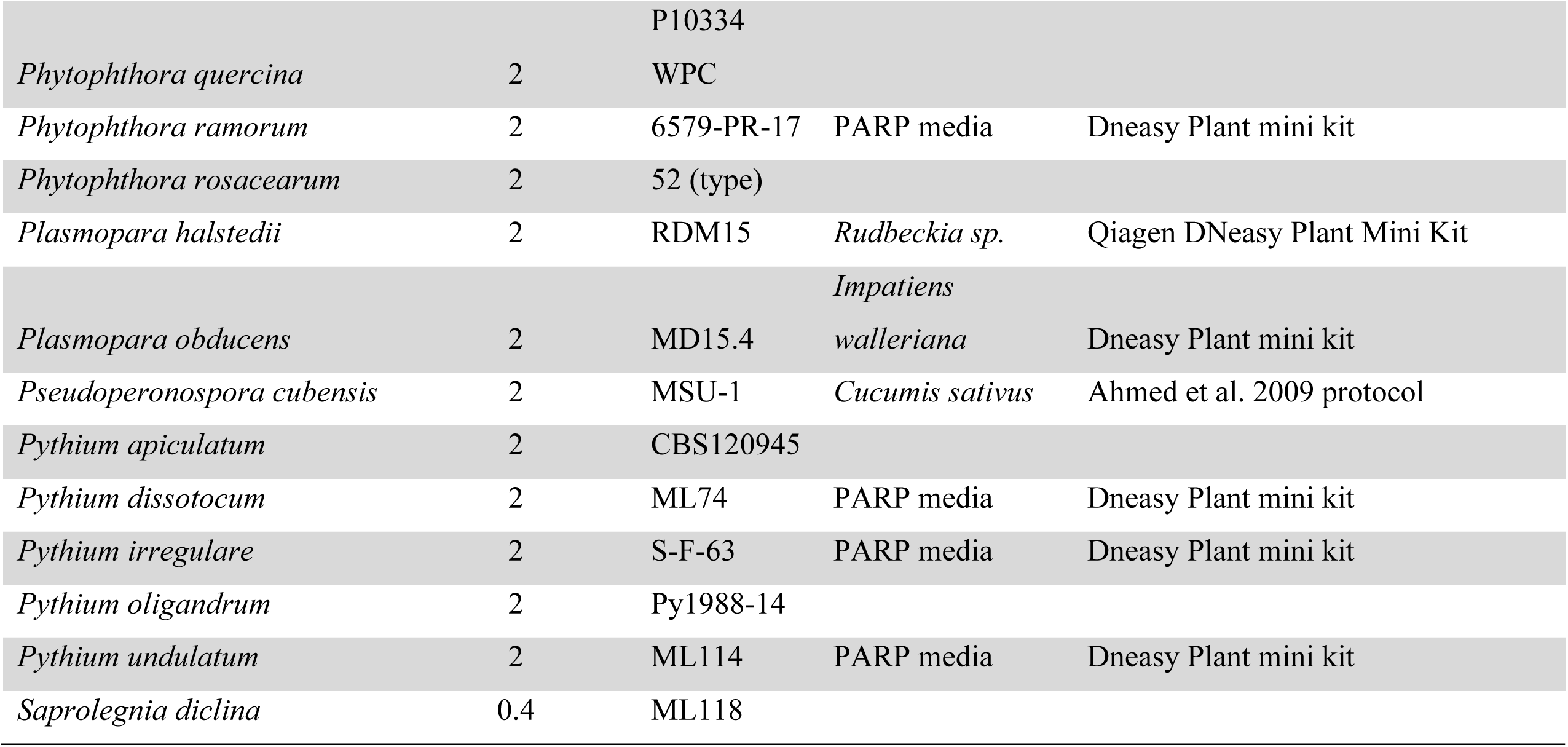
List of species included in the mock community. Table includes final DNA concentration in the mock community, isolate name, growth medium, and DNA extraction method.

| Species | Concentration |  | Isolate | Growth Medium | Extraction Method |
| --- | --- | --- | --- | --- | --- |
|  | (ng/μL) |  |  |  |  |
| *Aphanomyces euteiches* | 2 |  | MER4 | CMA | MPBio FastDNA |
| *Peronospora effusa* | 2 |  | n/a | *Spinacia oleracea* | Macherey-Nagel NucleoSpin Plant II |
| *Peronospora schachtii* | 2 |  | n/a | *Spinacia oleracea* | Macherey-Nagel NucleoSpin Plant II |
|  |  |  | CBS119171 |  |  |
| *Phytopythium citrinum* | 2 |  | (Type) |  |  |
| *Phytophthora cinnamomi* | 2 |  | Cin-TJ-130 | PARP media | Dneasy Plant mini kit |
| *Phytophthora citrophthora* | 2 |  | 6400-PR-16 | PARP media | Dneasy Plant mini kit |
| *Phytophthora himalsilva* | 2 |  | 531-P-13 | PARP media | Dneasy Plant mini kit |
| *Phytophthora hydrogena* | 2 |  | 46A3 (type) |  |  |
| *Phytophthora infestans* | 2 |  | 1826-PI-07 |  |  |
| *Phytophthora ipomoeae* | 2 |  | 1990-PI-07 |  |  |
| *Phytophthora kernoviae* | 2 |  | 55-PK-05 | PARP media | Dneasy Plant mini kit |
|  |  |  | 14Mar- |  |  |
| *Phytophthora pluvialis* | 0.7 |  | EMR | PARP media | Dneasy Plant mini kit |

|  |  |  |  |  |
| --- | --- | --- | --- | --- |
|  |  | P10334 |  |  |
| *Phytophthora quercina* | 2 | WPC |  |  |
| *Phytophthora ramorum* | 2 | 6579-PR-17 | PARP media | Dneasy Plant mini kit |
| *Phytophthora rosacearum* | 2 | 52 (type) |  |  |
| *Plasmopara halstedii* | 2 | RDM15 | *Rudbeckia sp.* | Qiagen DNeasy Plant Mini Kit |
|  |  |  | *Impatiens* |  |
| *Plasmopara obducens* | 2 | MD15.4 | *walleriana* | Dneasy Plant mini kit |
| *Pseudoperonospora cubensis* | 2 | MSU-1 | *Cucumis sativus* | Ahmed et al. 2009 protocol |
| *Pythium apiculatum* | 2 | CBS120945 |  |  |
| *Pythium dissotocum* | 2 | ML74 | PARP media | Dneasy Plant mini kit |
| *Pythium irregulare* | 2 | S-F-63 | PARP media | Dneasy Plant mini kit |
| *Pythium oligandrum* | 2 | Py1988-14 |  |  |
| *Pythium undulatum* | 2 | ML114 | PARP media | Dneasy Plant mini kit |
| *Saprolegnia diclina* | 0.4 | ML118 |  |  |

## Notes

http://www.oomycetedb.org

https://github.com/grunwaldlab/OomyceteDB

### Competing Interest Statement

The authors have declared no competing interest.

